# Engineering Plant Architecture via CRISPR/Cas9-mediated Alteration of Strigolactone Biosynthesis

**DOI:** 10.1101/254698

**Authors:** Haroon Butt, Muhammad Jamil, Jian You Wang, Salim Al-Babili, Magdy Mahfouz

## Abstract

Precision plant genome engineering holds much promise for targeted improvement of crop traits via unprecedented single-base level control over the genetic material. Strigolactones (SLs) are a key determinant of plant architecture, known for their role in inhibiting shoot branching (tillering). Here, we used CRISPR/Cas9 in rice *(Oryza sativa)* for targeted disruption of *CAROTENOID CLEAVAGE DIOXYGENASE 7 (CCD7),* which controls a key step in SL biosynthesis. The *ccd7* mutants exhibited a striking increase in tillering, combined with a dwarf phenotype, which could be rescued by application of the synthetic SL analog GR24. *Striga* germination assays and liquid chromatography–mass spectrometry analysis showed that root exudates of *ccd7* mutants were also SL deficient. Taken together, our results show the power of CRISPR/Cas9 for targeted engineering of plant architecture and for elucidating the molecular underpinnings of architecture-related traits.

## Introduction

Technologies that facilitate efficient, robust, and precise engineering of the plant genome can be used for targeted improvement of crop traits, and will pave the way for increasing plant yield and improving food security (Moshelion and Altman 2015). Plant architecture is dynamically regulated by developmental and environmental factors and has key effects on yield. For example, in the Green Revolution, random mutagenesis and harnessing of natural variants of key architecture genes in grain crops yielded varieties with shorter heights and more branches, resulting in significant improvement of crop productivity (Sakamoto and Matsuoka 2004).

Genome engineering requires molecular scissors capable of making precise double strand breaks (DSBs) in the genome (Sander and Joung 2014; Voytas and Gao 2014). Such DSBs are repaired by either the imprecise non-homologous end joining (NHEJ) repair or the precise homology directed repair (HDR) pathways (Symington and Gautier 2011; Voytas and Gao 2014; Butt et al. 2017). Harnessing the cellular repair pathways of the DSBs, a variety of user-desired genetic outcomes can be generated. Different platforms of site-specific nucleases (SSNs) have been used to engineer the eukaryotic genomes including homing endonucleases, zinc finger nucleases (ZFNs), transcription activator-like effector nucleases (TALENs) (Kim and Kim 2014). A novel class of SSNs, clustered regularly interspaced palindromic repeats (CRISPR)/CRISPR associated (Cas) 9 system is profoundly revolutionizing our ability to engineer the plant genome (Ran et al. 2013; Doudna and Charpentier 2014; Hsu et al. 2014; Sander and Joung 2014; Eid et al. 2016; Eid and Mahfouz 2016; Zaidi et al. 2016). Recently, different CRISPR/Cas systems have been harnessed to edit and regulate the RNA, and for other RNA manipulations (Mahas et al. 2018; Aman et al. 2018).

Strigolactones are a novel class of plant hormones that play an essential role in establishing plant architecture, determining the number of shoot branches/tillers and regulating the growth of primary and lateral roots (Gomez-Roldan et al. 2008; Umehara et al. 2008; Waldie et al. 2014; Al-Babili and Bouwmeester 2015; Jia et al. 2017). SLs also participate in biotic and abiotic stress responses (Ha et al. 2014; Torres-Vera et al. 2014; Decker et al. 2017). Furthermore, plant roots release SLs into the rhizosphere to trigger hyphal branching in mycorrhizal fungi for establishment of the beneficial arbuscular mycorrhizal symbiosis used by around 80% of land plants to improve nutrient uptake (Gutjahr and Paszkowski 2013; Bonfante and Genre 2015). However, seeds of root-parasitic weeds of the genus *Striga* perceive SLs as a germination signal ensuring the presence of a nearby host (Xie et al. 2010). Infestation by *Striga hermonthica* and related parasitic plants causes enormous yield losses in many crops, such as cereals and different *Solanaceae* species, representing a severe problem for agriculture in sub-Saharan Africa, Southern Europe, the Middle East and Asia (Parker 2009).

Analysis of SL-deficient and SL-perception mutants paved the way for the elucidation of the major steps in SL biosynthesis and signaling (Al-Babili and Bouwmeester 2015; Waters et al. 2017; Jia et al. 2017). For example, the increased branching/tillering phenotype of *carotenoid cleavage dioxygenase 7 (ccd7)* and *ccd8* mutants from *Arabidopsis thaliana,* pea *(Pisum sativum),* petunia *(Petunia hybrida),* and rice suggested the role of these enzymes in the biosynthesis of a shoot branching inhibitor that was later identified as SL (Ruyter-Spira et al. 2013; Al-Babili and Bouwmeester 2015). SLs are carotenoid-derivatives synthesized from all-*trans*-β-carotene via a pathway involving the all-*trans*/9-*cis*-β-carotene isomerase (DWARF27 in rice) that forms 9-*cis*-β-carotene (Alder et al. 2012; Bruno and Al-Babili 2016). In the next step, the stereospecific CCD7 cleaves 9-*cis*-β-carotene into the volatile β-ionone and a 9-*cis*-β-apo-10’-carotenal (Alder et al. 2012; Bruno et al. 2014). This *cis*-configured intermediate is the substrate of CCD8 that catalyzes a combination of reactions, including repeated deoxygenation and intramolecular rearrangements, which yield carlactone and a C_8_-product (ω-OH-(4-CH3) heptanal) (Alder et al. 2012; Bruno et al. 2017). Carlactone is the precursor of canonical and non-canonical SLs (Jia et al. 2017). In Arabidopsis, carlactone is converted by a cytochrome P450 of the 711 clade (MAX1) into carlactonoic acid, followed by methylation by an unknown enzyme and hydroxylation by lateral branching oxidoreductase into a yet unidentified SL (Abe et al. 2014; Brewer et al. 2016). Rice MAX1 homologs convert carlactone into the known SLs 4-deoxyorobanchol and orobanchol, likely via carlactonoic acid (Zhang et al. 2014; Iseki et al. 2017).

The first step in SL signal transduction is the binding of SL to the receptor, DWARF14 (D14) in rice, an α/β-hydrolase that hydrolyzes SL ligands and forms a covalent bond with one of the two hydrolysis products (D-Ring) (Hamiaux et al. 2012; de Saint Germain et al. 2016; Yao et al. 2016). These steps are accompanied by a conformational change that enables the interaction with an F-box protein (MAX2 in Arabidopsis; D3 in rice), which is a constituent of an SKP1-CUL1-F-box-protein (SCF)-type ubiquitin ligase complex, initiating the 26S proteasomal degradation of target transcription repressors (Jiang et al. 2013; Zhou et al. 2013; Waldie et al. 2014; Soundappan et al. 2015).

Targeted mutation of key enzymes in SL biosynthesis or perception can be used to engineer plant architecture in ways that improve yield. Moreover, in contrast to traditional, time-consuming approaches involving breeding natural alleles into elite varieties, genome editing provides a targeted, precise and rapid method to improve key plant traits in agronomically relevant, locally adapted varieties. OsCCD7 catalyzes a key step in SL biosynthesis and an *OsCCD7* point mutation (C-T) in the Nanjing 6 background, *htd1 (high-tillering and dwarf 1),* causes a dwarf phenotype and production of a high number of tillers (Zou et al. 2006; Zhang et al. 2011). This indicates the possibility of manipulating plant height and tillering, key agronomic traits, by genome editing of *OsCCD7.* In this work, we report the engineering of SL biosynthesis by CRISPR/Cas9-mediated mutation of *OsCCD7.* Our data provide a proof-of-principle for translational application of knowledge on SL biosynthesis for improvement of plant architecture.

## Results

### Targeted engineering of *CCD7* mutants in rice via CRISPR/Cas9

We used CRISPR/Cas9 for targeted mutagenesis of *OsCCD7* to generate variants for translational research and to enhance our understanding of the diverse functions of this protein. The *OsCCD7* gene *(LOC_Os04g46470)* has 7 exons encodeing a protein of 609 amino acids (Fig. 1B) that mediates a key step in SL biosynthesis (Fig. 1A). For targeted mutagenesis of *OsCCD7,* we engineered two gRNAs, gRNA1 (ACCTACTACCTCGCCGGGCCGGG) and gRNA2 (AAGAACCTCACTTTTCCAATGGG) targeting the 1^st^ and the 7^th^ exon, respectively (Fig. 1B). We expressed the gRNAs under the control of the RNA polymerase III *U3* promoter *(U3::gRNA)* and with a poly-T termination signal. We cloned this into a binary vector containing the T-DNA region with the Cas9 endonuclease driven by the *OsUbiquitin* promoter for *in planta* expression. We transferred the resulting vector into *Agrobacterium tumefaciens* strain EHA105 and transformed rice calli, as previously reported (Butt et al. 2017).

**Fig. 1:**
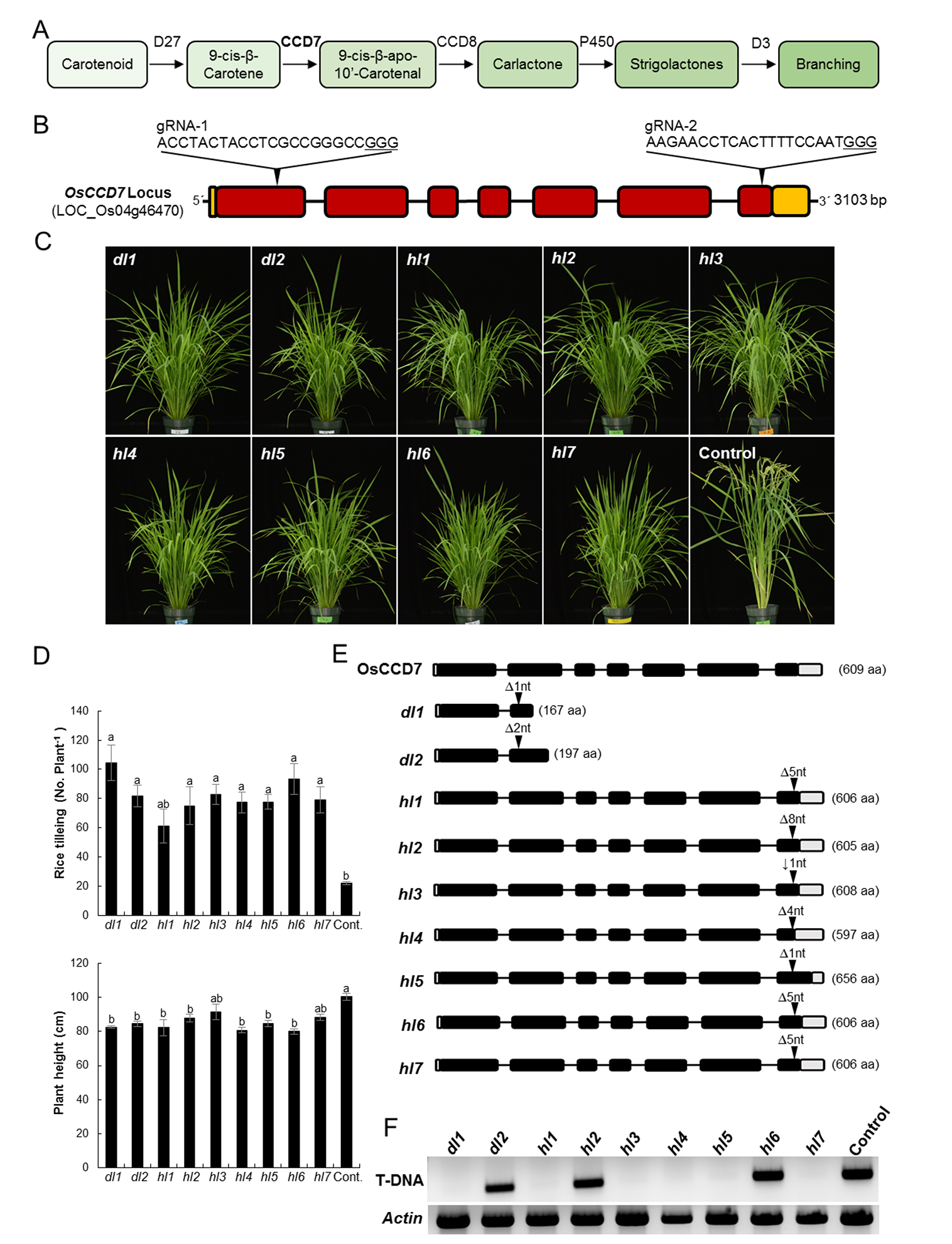
*OsCCD7* mutagenesis produced high-tillering and dwarf phenotypes: Schematic of the strigolactone biosynthesis pathway showing branching/tillering inhibition activity. Carotenoid Cleavage Dioxygenase 7 (CCD7) catalyzes 9’-*cis*-β-carotene at the initial steps of SL biosynthesis. **(B)** Gene model of *OsCCD7/ D17* (Dwarf 17)/ *HTD1* (High-Tillering Dwarf 1), also called MAX3 ortholog (LOC_Os04g46470). Two gRNA were used to target the *OsCCD7* locus. gRNA-1 was designed to target the 1^st^ exon to produce a mutation similar to *d17.* The nomenclature used for these mutant lines is *dl (d17-like).* Two T1 lines were produced, *dl1* and *dl2.* gRNA-2 was designed to target the 7^th^ exon. The mutant lines produced were similar to *htd1* and these lines were named as *hl (htd1-like).* Seven independent lines were produced, *hl1* to *hl7.* (**C** and **D**) High-tillering and dwarf phenotypes were observed for mutant plants. Tillering and plant height were recorded for all genotypes at eight weeks after germination. All of the genotypes showed significant increases in tillers per plant and decreased plant height compared to control. T_2_ generation of mutants were genotyped and mono-allelic lines were identified. Each of these mutations produced a protein variant. The nucleotide indels and protein alignments are shown in supplementary figure 2 and supplementary figure 3 respectively. **(F)** Analysis of the T_3_ generation showed some of the mutant lines do not carry a T-DNA. T-DNA-specific PCR analysis indicated that *dl1, hl1, hl3, hl4, hl5* and *hl5* are non-transgenic mutated plants. *Actin* PCR was done as a control.

After regeneration of T1plants from the calli, we examined the resulting plants for mutations in *OsCCD7.* We recovered 22 T1 transgenic plants corresponding to sgRNA1 and 17 T1 transgenic plants corresponding to sgRNA2 and genotyped these plants by PCR amplifying the region encompassing the target site of the sgRNA. PCR amplicons were cloned and sequenced. For gRNA1, 8/22 plants showed bi-allelic mutations of the target site with formation of insertion/deletion mutations (indels) including deletions of 1–27 bp. Similarly, for gRNA2, all of the plants were bi-allelic with indels. Some plants, however, exhibited only monoallelic mutations. These data show the high efficiency of targeted mutagenesis for *CCD7* by gRNAs targeting the 1^st^ and 7^th^ exons. We used the nomenclature *d17-like (dl)* for mutants produced by gRNA1. Two mono-allelic mutants, *dl1* and *dl2* with one‐ and two-bp deletions, respectively, were used for further studies (Supplementary Fig. 1A). For gRNA2 mutants, we used the nomenclature *htd1-l^i^ke (hl).* Seven bi-allelic mutants *hl1–hl7* were used for further studies (Supplementary Fig. 1A).

### The *ccd7*mutants exhibit increased tillering and dwarf phenotypes

Our genotyping data revealed the presence of several *ccd7* mutants resulting in complete or partial functional knockout phenotypes. Phenotyping the *T1* plants can accelerate functional analysis, but the presence of two alleles can complicate interpretation. To examine this, we phenotyped the T1 plants for the number of tillers and plant height (Supplementary Fig. 1B, 1C). Our data indicate that the mutants produced increased numbers of tillers and reduced plant height, reminiscent of the *ccd7* mutants isolated by conventional methods in other plant species (Supplementary Fig. 1B, 1C). *hl5* and *hl2* showed the highest numbers of tillers per plant, while dl2 and *hl1* exhibited the most pronounced dwarf phenotype among all mutants (Supplementary Fig. 1C). However, *hl4* showed high tillering accompanied with less reduced height, (Supplementary Fig. 1C).

Most of the *hl* mutations observed were bi-allelic in the T1 generation. To analyze and correlate the phenotypes with a particular protein variant, T_2_ plants were genotyped by sequencing. Mono-allelic, homozygous mutant plants were identified and used for further studies. All of the mutant lines, including *hl4,* showed pronounced high tillering and dwarf phenotypes (Fig. 1C, 1D). Among these mutant lines, *dl1* and *hl6* showed the highest number of tillers per plant (Fig. 1D). Each of the mutant lines harbors a particular mutation that leads to a CCD7 protein variant (Fig. 1E, Supplementary Fig. 2). These protein variants produce variation in tillering and plant height (Fig. 1D).

One advantage of CRISPR/Cas9 mutagenesis is that the mutation can be segregated away from the T-DNA expression construct used to produce Cas9 and the gRNA. To analyze whether these lines harbor a T-DNA, we examined the T_3_ generation of these mutants. We found some of the plants have no T-DNA but do have mutations in *CCD7* (Fig. 1F). This further showed that non-transgenic mutated plants of agricultural importance can be produced.

### *ccd7* mutants exhibit impaired SL biosynthesis

To further show that the *ccd7* mutant phenotype results from defects in SL biosynthesis, we tested whether treatment of the *ccd7* mutants with the synthetic SL analogue GR24 could restore their phenotypes to wild type. Progeny seeds were germinated on filter paper in Petri dishes and one-week-old seedlings were grown in 50-ml tubes for another week. The synthetic SL analogue GR24 was applied at 2.5 μM concentration for three weeks, then tiller numbers were counted. Application of GR24 to the *ccd7* mutants decreased tiller numbers to wild-type levels (Fig. 2A–C). The observed number of tillers of *ccd7* mutants was 7 tillers per plant on average in untreated (Mock) plants and this decreased to 1 tiller per plant with GR24 treatment (Fig. 2A-C).

**Fig. 2:**
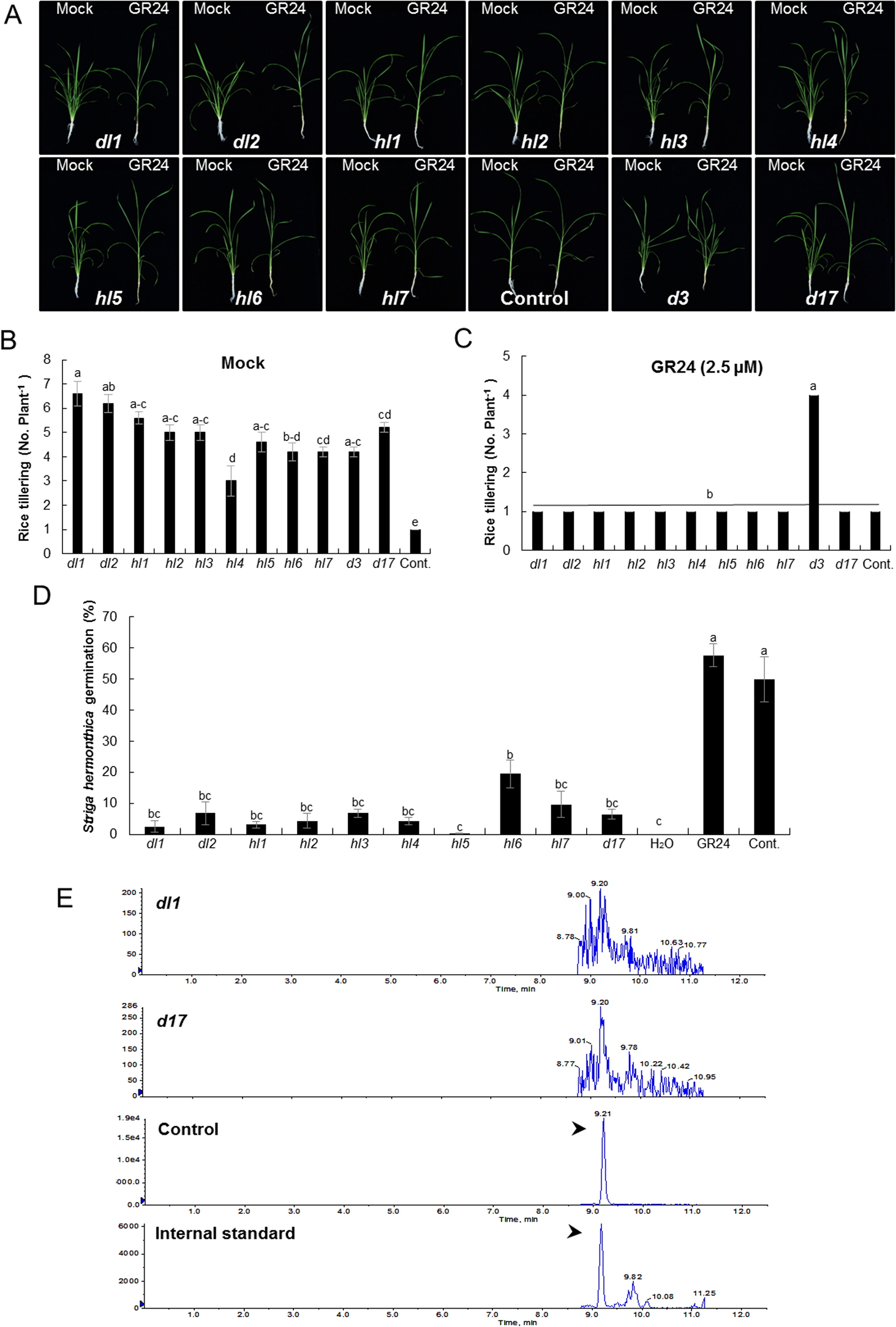
The *ccd7* mutants showed impaired SL biosynthesis and decreased SL-dependent biological activity: **(A)** GR24 treatment rescued the tillering phenotype of all T_2_-mutant lines. One-week-old seedlings were grown in 50-ml tubes and GR24, a synthetic SL analog, was applied twice a week at 2.5 μM for three weeks. Cas9ox (Ctrl), the SL-deficient mutant *d17,* and the SL-perception mutant *d3* were used as controls. Tillering inhibition by GR24 feeding indicated that the high-tillering phenotype appeared because of lack of SLs biosynthesis. **(B-C)** Tillers per plant were recorded with or without GR24 treatment after three weeks of application. No. of tillers per plant were significantly reduced after GR24 treatment. Bars represent means ±SE (*n*=6). Means not sharing a letter in common differ significantly at *P*_0_.05. **(D)** *Striga* germination bioassay was conducted to measure SL bioactivity in T_2_ mutant lines. The rice plants were grown under normal conditions for 3 weeks. Then in 4^th^ week each line was grown under phosphate-deficient conditions for another week. The SLs were extracted from root exudates of each line and applied to pre-conditioned *Striga* seeds. Very low *Striga* seed germination was observed in mutants compared to control, indicating SL deficiency in these lines. Graph shows percent *Striga* germination in response to root exudates collected from each line. Cas9ox (WT) and d17 were used as control plant lines; in addition, the SL analog GR24 and H_2_O were applied to *Striga* seeds with no root extracts for controls to show maximum and minimum germination. **(E)** LC-MS∕MS analysis using multiple reaction monitoring (MRM) of rice root exudates. The MRM transitions for various SLs from root exudates of control, *d17and ccd7* T_2_-mutant lines were observed. All of the mutants and *d17did* not show any detectable signals for SLs. The chromatogram showed peaks for 4-deoxystrigol in the internal standard and control. One of the mutants *(dl1)* chromatogram is shown as an example.

To confirm the SL deficiency of the *ccd7* mutants, we quantified the levels of SLs in the corresponding root exudates, using liquid chromatography quadruple time-of-flight tandem mass spectrometry (LC-MS/MS). The LC-MS/MS data indicated that none of the mutants produce detectable levels of SLs (Fig. 2E).

### The *ccd7* mutants affected *Striga* germination

SLs induce the germination of parasitic seeds of the *Orobancheaceae* family, including *Striga;* therefore, we measured the ability of exudates from *ccd7* mutant roots to stimulate the germination of *Striga* seeds. We found a significant reduction in *Striga* germination with the root exudates of *ccd7* mutants as compared to wild type (Fig. 2D). The different *ccd7* mutants exhibited different levels of SL production and different plant architecture phenotypes. However, the reduction of *Striga* germination was not correlated with the SL levels in the *ccd7* mutants measured by LC-MS. This might be due to higher sensitivity of the *Striga* bioassay, as compared to LC-MS analysis.

## Discussion

Use of the CRISPR/Cas9 system to precisely manipulate the plant genome in a user-defined manner opens myriad possibilities for translational applications in agriculture (Voytas and Gao 2014; Schiml and Puchta 2016; Eid and Mahfouz 2016; Zaidi et al. 2016; Aman et al. 2018). Genes controlling plant architecture are prime targets for engineering plant varieties with higher yields; however, to avoid unintended effects, such engineering requires a strong understanding of the mechanisms by which such genes function. In recent years, multiple lines of inquiry have provided a substantial understanding on the biology of SLs, including their biosynthesis, transport, signaling, and interactions with other hormonal responses (Seto and Yamaguchi 2014; Jia et al. 2016; Waters et al. 2017). SLs control shoot branching and stimulate the germination of parasitic plants; therefore, the manipulation of SL biosynthesis may reduce crop losses and increase yield (Lopez-Raez et al. 2009; Conn et al. 2015). For example, reduction of SLs in plants and root exudates could compromise the germination of parasitic plant seeds, and thus reduce crop losses. Indeed, reduction of SL production has shown promise in plant species including rice, pea, fava bean *(Vicia faba),* tomato *(Solanum lycopersicum),* and maize *(Zea mays).* However, significant reductions in SL levels may compromise other factors, such as the formation of arbuscular mycorrhizal symbioses, and thereby ultimately prove to be a counterproductive strategy. Moreover, finding germplasm of a given crop species with reduced and fine-tuned SL production remains challenging due to the limited genetic diversity of crop varieties. The CRISPR/Cas9 system can be used to produce the much-needed genetic diversity and generate varieties with fine-tuned levels of proteins or metabolites such as SLs.

Here, we employed the CRISPR/Cas9 system for targeted engineering of *CCD7* to produce rice varieties with reduced SL biosynthesis. We engineered the CRISPR/Cas9 system to target two exons in *CCD7* with the aim of producing complete and partial knockout phenotypes. Having rice varieties with reduced levels of CCD7 may help to fine-tune the levels of SLs within the plant and the root exudates. We also generated non-transgenic and mono-allelic *ccd7* mutants. Phenotypic analysis of tillering and plant height indicated the diversity among these mutant lines. *Striga* germination bioassays also showed similar variation among mutants. The *hl6* mutant showed the highest variability, producing the most tillers per plant, but also giving highest *Striga* germination (Fig. 1D, 2D).

Our study provides an example of the application of CRISPR/Cas9 to manipulate SL biosynthesis and production for targeted translational applications. Future studies would focus on producing varieties of rice and other plants with fine-tuned SL levels to improve plant architecture and reduce the germination of parasitic plants without compromising other agronomic traits. For example, such approaches could involve generation of protein variants of the SL signaling components that are very sensitive to endogenous SL levels, thereby maintaining SL responses and mediating feedback inhibition of SL biosynthesis, leading to proper fine-tuning of SL levels. Furthermore, targeted manipulation of the upstream transcriptional regulators of SL biosynthesis would provide another avenue to fine-tune SL levels in the target plant species for trait improvement. Other avenues include the production of enzymatic variants that produce less-stable SLs or produced different SL types in the target species. For example, different SLs might differentially affect tillering, plant stature, *Striga* germination, and establishment of arbuscular mycorrhizal symbioses; production of SLs with minimal activity for *Striga* germination, but maximal activity for beneficial symbiotic fungi might improve yields and reduce losses. In conclusion, engineering the plant genome via CRISPR/Cas9, and other developing systems, unlocks myriad possibilities to fine-tune SL biosynthesis and responses for targeted improvement of crop traits.

## Materials and Methods

### Plant materials and vector construction

*Oryza sativa* L. ssp. *japonica* cv Nipponbare was used for all experiments. The pRGEB32 vector was used for callus transformations (Xie et al., 2015(Butt et al. 2017)). The expression of Cas9 was driven by *OsUbiquitin* and the gRNA was expressed as a polycistronic tRNA-gRNA under the *OsU3* promoter. Two gRNAs were designed and transformed independently. gRNA-1 was designed to target the genomic sequence 289 to 308 bp (5’-ACCTACTACCTCGCCGGGCCGGG-3’); the underlined <u>GGG</u> represents the PAM sequence. gRNA-2 was designed to target the genomic sequence 2416 to 2435 bp (5’-AAGAACCTCACTTTTCCAAT<u>GGG</u>-3’); the underlined <u>GGG</u> represents the PAM sequence. The pRGEB32 vector transformed without gRNA was transformed into Nipponbare and used as the control. *d3-1* and *d10-1,* in the Shiokari background (Ishikawa et al. 2005) were used as tillering dwarf mutants.

### Rice transformation and mutant screening

*Agrobacterium-*mediated rice transformation was performed as described previously (Butt et al., 2017). Transgenic rice plants were grown in a greenhouse at 28 °C. After one week when plants were established on soil, DNA was extracted from a leaf sample. PCR was done using gene-specific primers. Purified PCR products were cloned using the CloneJET PCR Cloning Kit (K1231). Sanger sequencing was done for at least ten colonies to analyze the mutation.

### Rice tillering bioassays

Rice seeds were surface-sterilized with 2.5% sodium hypochlorite for 10 min. The seeds were then washed thoroughly with sterile MilliQ water and imbibed in water for 2 days at 30*°C* in the dark. The pre-germinated seeds were shifted to 90-mm Petri dishes on a filter paper moistened with 5 ml ½-strength Murashige and Skoog (Gilbert et al.) medium. The sealed plates were kept at 30*°C* for one more night to develop small seedlings. The plates with small seedlings were kept under fluorescent white light (130–180 μM m^-2^ s^-1^) for 7 days. One-week-old uniform seedlings were selected to grow in a microfuge tube, fixed on top of 50-ml tubes (one seedling per tube) filled with modified half-strength Hoagland nutrient solution. After one week, the rice plants were supplemented with GR24 at 2.5 μM. The GR24 was added six times, twice a week. Number of tillers per plant and plant height of all lines including control, *d3,* and d17 were measured at final harvest.

### Quantitation of SLs from rice root exudates

Rice seeds were surface sterilized with 2.5% sodium hypochlorite and 10 μl Tween-20 and germinated on ^½^-strength MS agar at 30 *°C* under fluorescent white light (130-180 μM m^-2^ s^-1^) to establish seedlings. Seven-day-old uniform seedlings were transferred to 50-ml tubes (three seedlings per tube) containing modified half-strength Hoagland nutrient solution and grown in a growth cabinet with normal phosphorus (P) supply for one week. Then, rice seedlings were subjected to P deficiency for another week. On the day of root exudate collection, rice seedlings were first refreshed with P-deficient Hoagland nutrient solution for 6 h and then root exudates were collected from each tube. SLs were collected from each root exudate sample for LC/MS-MS analysis and *Striga* bioassays. The SPE C18 column (Grace Pure) was pre-conditioned by solvation (6 ml of methanol) and an equilibration (6 ml of water). The internal standard D_6_-5-Deoxystrigol (0.672 ng) was added in each 50 ml root exudate. The root exudates were run through the preconditioned SPE C_18_ column. After washing with 6 ml of water, SLs were eluted with 5 ml of acetone. The SL fraction (acetone-water solution) was concentrated to SL aqueous solution (∼1 ml), followed by extraction with 1 ml of ethyl acetate. Then 750 μl of SL-enriched organic phase was transferred to a 1.5-ml tube and evaporated to dryness. The sample was re-dissolved in 100 μl of acetonitrile:water (25:75, v:v) and filtered through a 0.22-μm filter for LC-MS/MS analysis. SLs were analyzed using HPLC-Q-Trap-MS/MS with MRM mode. Chromatographic separation was achieved on an Acquity UPLC BEH C18 column (50 × 2.1 mm; 1.7 μm; Waters). Mobile phases consisted of water:acetonitrile (95:5, v:v) and acetonitrile, both containing 0.1% formic acid.

### *Striga hermonthica* bioassays

The *Striga hermonthica* seed germination bioassay was conducted as described previously (Jamil et al., 2012). The *Striga* seeds were preconditioned for 10 days at 30°C under moist conditions. The pre-conditioned *Striga* seeds were supplied with 50 μL acetone-free SL, collected from the root exudates for each mutant line as described above. Wild type and the *d17* mutant were included as positive and negative controls, respectively. After SL application, *Striga* seeds were incubated at 30*°C* in the dark for two days. Germinated (seeds with radicle) and non-germinated seeds were counted under a binocular microscope to calculate germination rate

## Acknowledgements

We would like to thank members of the laboratory for genome engineering at KAUST for their helpful discussions and critical reading of the manuscript. This study was supported by King Abdullah University of Science and Technology.

